# LD-CNV: rapid and simple discovery of chromosomal translocations using linkage disequilibrium between copy number variable loci

**DOI:** 10.1101/2021.06.18.449059

**Authors:** Luca Comai, Kirk Amundson, Benny Ordonez, Xin Zhao, Guilherme Tomaz Braz, Jiming Jiang, Isabelle Henry

## Abstract

Large scale structural variations, such as chromosomal translocations, can have profound effects on fitness and phenotype, but are difficult to identify and characterize. Here, we describe a simple and effective method aimed at identifying translocations using only the dosage of sequence reads mapped on the reference genome. We binned reads on genomic segments sized according to sequencing coverage and identified instances when copy number segregated in populations. For each dosage-polymorphic 1Mb bin, we tested linkage disequilibrium (LD) with other variable bins. In nine potato (*Solanum tuberosum*) dihaploid families translocations affecting pericentromeric regions were common and in two cases were due to genomic misassembly. In two populations, we found evidence for translocation affecting euchromatic arms. In cv. PI 310467, a non-reciprocal translocation between chromosome 7 and 8 resulted in a 5-3 copy number change affecting several Mb at the respective chromosome tips. In cv. “Alca Tarma”, the terminal arm of chromosome 4 translocated to the tip of chromosome 1. Using oligonucleotide-based fluorescent in situ hybridization painting probes (oligo-FISH), we tested and confirmed the predicted arrangement in PI 310467. In 192 natural accessions of *Arabidopsis thaliana*, dosage haplotypes tended to vary continuously and resulted in higher noise, while LD between pericentromeric regions suggested the effect of repeats. This method should be useful in species where translocations are suspected because it tests linkage without the need for genotyping.

## Introduction

Genomic structural variation (SV) is common within plant populations (Swanson-Wagner *et al*. 2010) and has a profound effect on the phenotype of individuals (Díaz *et al*. 2012; Maron *et al*. 2013; Bastiaanse *et al*. 2019). SV is manifested at multiple scales, from single genes to large chromosomal regions, such as variable heterochromatic blocks in the pericentromeres of potato (*Solanum tuberosum*) (Gong *et al*. 2012; Zhang *et al*. 2014; de Boer *et al*. 2015; Hardigan *et al*. 2016). Small scale SV can alter gene structure and copy number. Large scale translocations and inversions can alter copy number, recombination, meiotic anaphase patterns, viability of gametes, gene structure, and evolutionary potential (Khush 1973; Rieseberg 2001). Translocations, which tend to be underdominant, i.e. to confer a heterozygous disadvantage (Rieseberg 2001), can persist in nature when they involve reciprocal exchanges of chromosome arms, resulting in copy-neutral, monocentric recombinant chromosomes. Nonreciprocal translocations are often deleterious because they can involve duplication and deletion of segments of the recombined chromosomes. However, in polyploids their effect is lessened by genomic redundancy.

Detection of translocation is not simple and often requires multiple approaches. SV loci display linkage disequilibrium (LD) with linked regions (Hinds *et al*. 2006; Locke *et al*. 2006). Translocations were originally detected by observing unusual lethality and unexpected linkage in Drosophila (Bridges 1923) and in plants (Belling and Blakeslee 1924). Later, they were confirmed cytologically (Belling and Blakeslee 1924; Muller 1929; McClintock 1930; Burnham 1956). The development of fluorescent in situ hybridization (FISH) facilitated their identification because each chromosome in a spread could be identified by cytological markers or by chromosome painting (He *et al*. 2018). Detection by FISH analysis requires chromosome-specific probes (Cremer *et al*. 1988), which can be readily developed using an oligonucleotide-based methodology (Zhang *et al*. 2021).

Genome sequencing can be informative of copy number changes and novel junctions (Alkan *et al*. 2011). Dosage analysis of sequence reads is informative when unbalanced translocations result in copy number variation (CNV), and when balanced translocations generate copy number variable progeny. CNV can be inferred by count of genomic sequencing reads. Regions with high or low read counts have likely undergone duplication or deletion. Read count, however, does not provide information on the context and position of copy number variable regions. Specifically, a duplication could involve transposition or be in tandem. The software CNVmap exploits heterozygous states arising from duplications to detect and map CNV (Falque *et al*. 2019). However, SNP between duplicated loci is a function of duplication age and may not be present in newly arisen structural variation. Evidence of a junction between two different chromosomes can be obtained by detecting sequencing reads that directly or indirectly span the junction. In practical terms, however, identification and characterization of a translocation is not simple, particularly in the absence of prior evidence pointing to its location. DNA analysis is complicated by the presence of many spurious signals in mapped sequenced reads, especially when using short sequencing reads, and by the frequent presence of repetitive DNA at translocation junctions (Chen *et al*. 2010). Long read technology can be effective at addressing these problems (Sedlazeck *et al*. 2018), although it is typically more expensive than short read sequencing.

Polyploid species are more likely to display large SV because their genome buffers the deleterious effects of copy number changes (Comai 2005). In polyploids, gametes carrying a large chromosomal deletion might still be viable because they carry a second normal copy of the affected chromosome. Cultivated potato is an autotetraploid, highly heterozygous organism that is clonally propagated, but capable of elevated outcrossing rates (Brown 1993). The potato genome displays a high density of nucleotide and structural polymorphism, suggesting high plasticity (Hardigan *et al*. 2016, 2017). From a cultivated tetraploid potato clone, sexual haploid progeny can be extracted by pollination with a haploid inducer (Fig.1-A). These haploids inherit a diploid genome consisting of the maternal gametic contribution and are therefore called dihaploids. Their diploid genome simplifies genetic analysis. Further, they are fertile when crossed to many wild diploid accessions, enabling the bridging of germplasm and introduction of valuable traits into cultivated potato (Peloquin *et al*. 1989; Rokka 2009).

**Figure 1.**
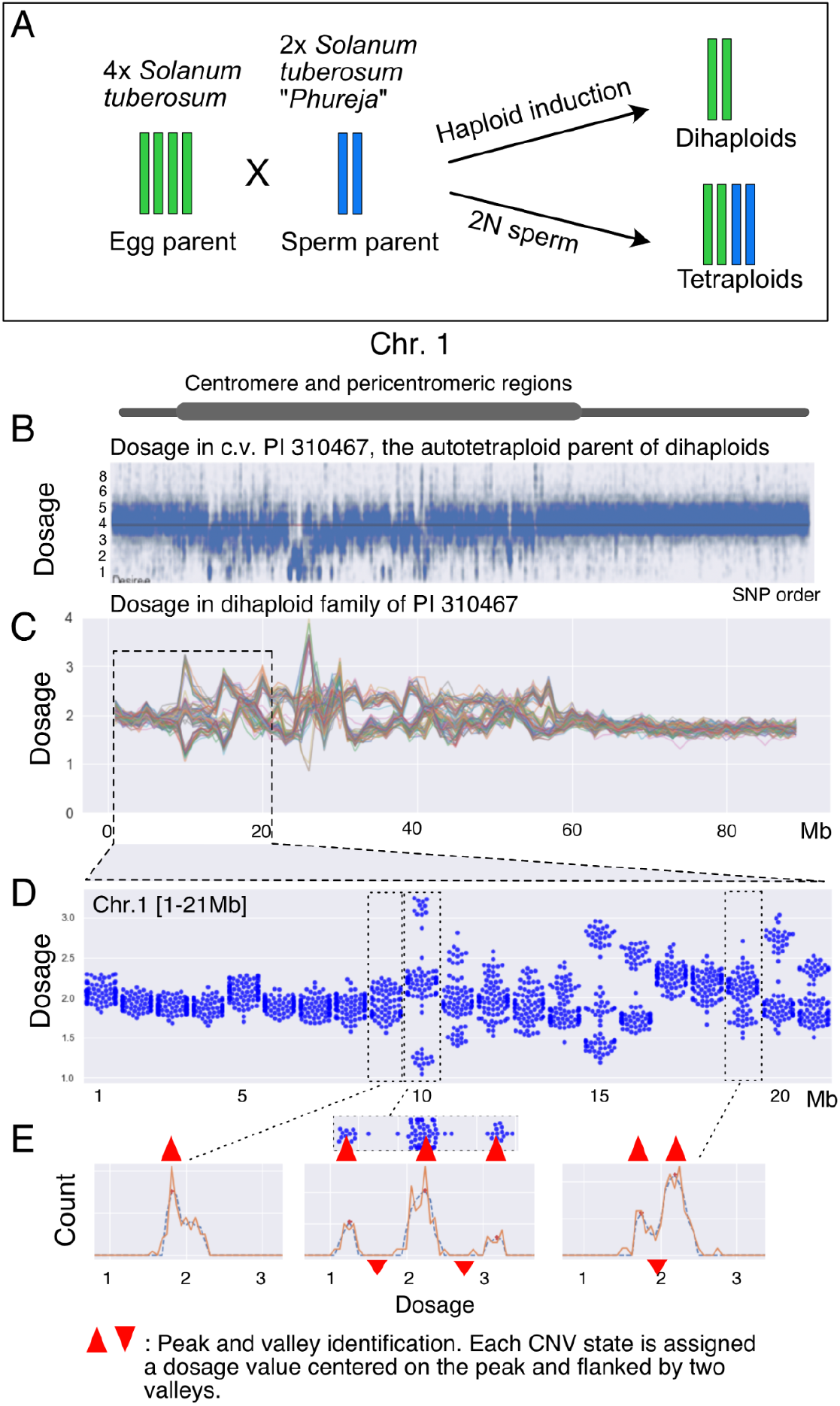
Dosage states of potato chromosome 1 in 84 dihaploid siblings. A. Haploid induction crosses generate progeny of different ploidy. Dihaploids inherit the genomic content of the egg gamete exclusively. Tetraploids are hybrids and receive contribution from both parental genomes. Triploid hybrids are also possible, but not depicted or analyzed here. B. Standardized coverage per SNP of chromosome 1 for the tetraploid parent PI 310467. C. Relative dosage states along chr.1 were derived by sequence coverage binned on 1Mb intervals and standardized on the maternal dosage. 84 maternal dihaploid individuals produced from the haploid induction cross *S*.*t*. PI 310467 x *S*.*t. group phureja* IvP48 are overplotted revealing recurring dosage trends and polymorphism, mainly associated with heterochromatic regions along the pericentromere. D. Swarm plots of dosage states in the first 21 Mb of chr.1. E. Cluster derivation by Peakutils. Peaks (upward red arrowheads) are identified by the algorithm, while valleys (downward red arrowheads) are defined as the mid distance between peaks. Dosage values flanked by two valleys were assigned to the corresponding clusters.

To investigate structural genomic variation, we developed a novel translocation detection approach and demonstrated its application in potato. Our method does not require genotyping or prior identification of polymorphism, such as SNPs or other common genetic markers. Instead, it leverages the reference genome sequence and dosage states inferred from sequencing the genome of related individuals, such as dihaploids of the same parent, in order to identify LD between structurally variant loci. The resolution of the identified structural variation scales with sequencing coverage, but even low coverage data can lead to interesting findings. Using several potato dihaploid populations, we demonstrate discovery of translocations in two families, as well as regions that could either be translocated and polymorphic, or assembly errors.

## Results

### Identification and LD analysis of CNV loci in a dihaploid population of potato

We generated a family of different ploidy types by crossing tetraploid potato variety PI 310467 to haploid inducer IvP48, producing a population called BB (Fig.1-A). PI 310467 is listed as Desiree in the GRIN germplasm collection but, although related, it differs from Desiree in its SNP profile (K.A., unpublished results). In the BB progeny, we selected and analyzed 84 2X haploids (dihaploids) by flow cytometry and low-pass genome sequencing. For comparison, we also analyzed 78 4X hybrid siblings. Mapped reads were counted in non-overlapping, 1Mb genomic intervals along the reference genome, and these counts were standardized to a mean of 2 using the corresponding maternal variety counts. When these standardized dosage values from all individuals were overplotted, CNV polymorphisms were readily visible, as displayed for chr.1 (Fig. 1, Suppl. Fig. 1). Invariant regions form unimodal distributions, while polymorphic regions form bimodal or multimodal distributions (Fig.1-BC). For each of these regions, individuals could be clustered based on their corresponding read counts (Fig.1-D, Suppl. Fig.1, Suppl. Fig.2). We used the Python utility Peakutils (Negri) to identify these clusters and assign each individual to a genomic dosage state, which can be viewed as a stepwise, quantitative phenotype resulting from additive alleles.

Consistent with previous reports (de Boer *et al*. 2015; Hardigan *et al*. 2016; Amundson *et al*. 2020a), dosage variation was common in pericentromeric regions (Fig.1-B, Suppl. Fig.1). To further characterize the associated structural changes, we asked whether dosage variation at different loci was independent. Linked loci, such as those in the same pericentromeric regions, would be expected to co-vary, i.e. to exhibit LD. Dosage states of unlinked loci should be independent. Unexpected LD suggests either epistasis or novel linkage, such as resulting from a translocation or genome assembly error. Therefore, for each SV locus, we tested the null hypothesis that its dosage states were independent of dosage states at another SV locus (Fig. 2).

**Figure 2.**
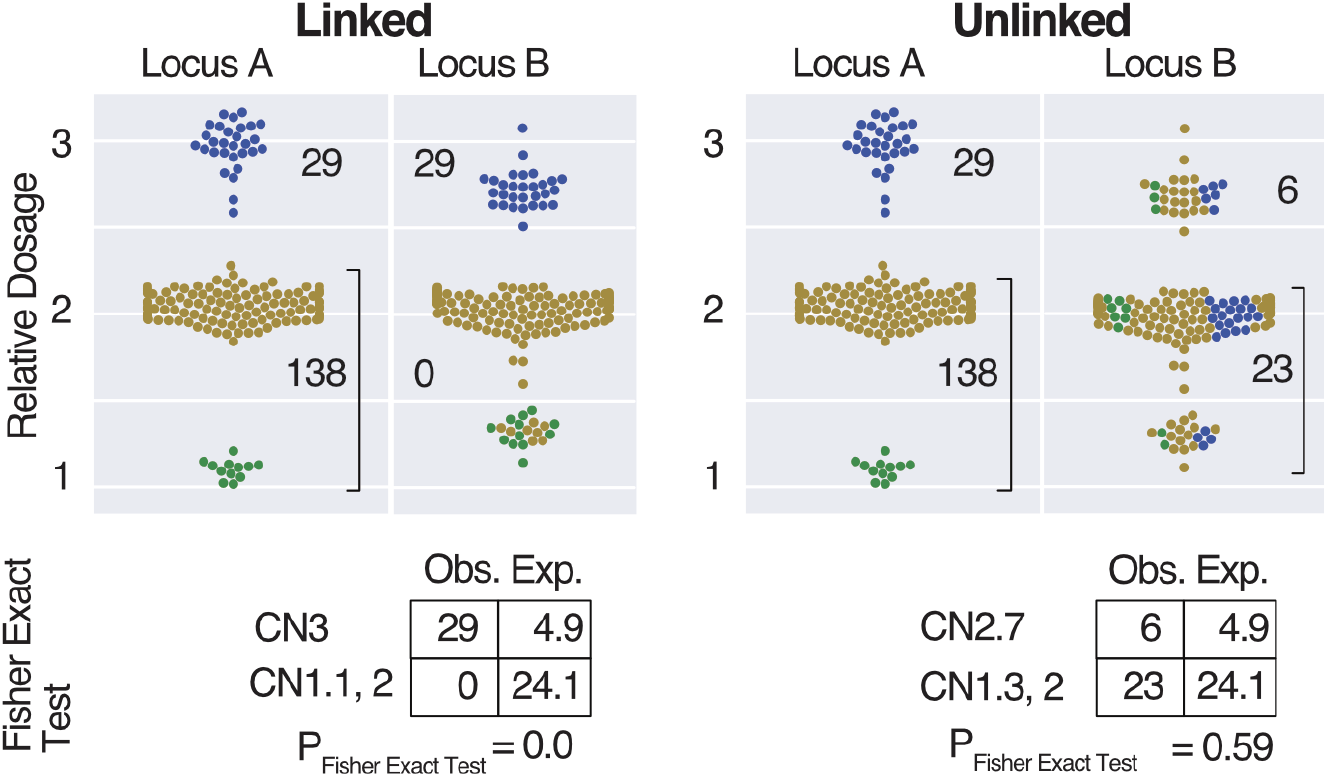
Testing random association between dosage states at different loci. Hypothetical loci A and B are illustrated for linked (left) and unlinked (right) states. The independence test compares dosage states at the two loci. Here, the top cluster is compared to the aggregate of the other two, producing a Fisher Exact 2×2 contingency table. Obs. = observed, Exp. = expected according to random association.

After correction for multiple testing (Benjamini and Hochberg 1995), associations between bins were plotted on a genomic matrix (Fig. 3). The potato genome was partitioned in 731 1Mb bins, of which 246 (33% of genome) displayed distinct dosage states. Most, 236, were in LD with at least another bin (corrected p < 0.05). This is not surprising, since LD is expected for physically linked loci. However, 65 bins were in LD with bins on other chromosomes, indicating that 26.7% of copy variable sequences display unexpected linkage. Loci in LD detected with FDR = 0.05 are marked in blue and black in the matrix (Fig. 3).

**Figure 3.**
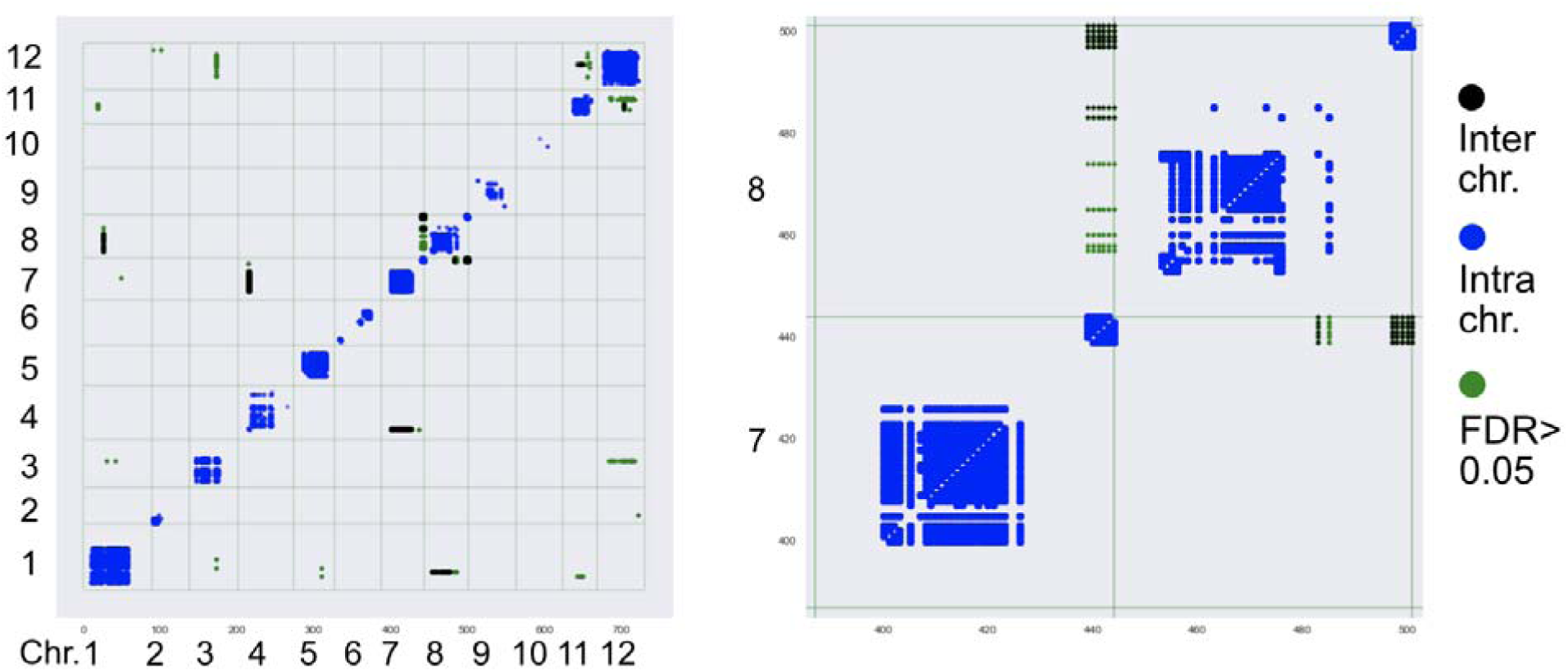
LD matrix of genomic dosage relationships. A twelve chromosome matrix on the left displays significant interactions (Fig. 2), with a zoomed view on two chromosomes on the right. Note that only CNV loci can be tested. To illustrate that most interactions are statistically significant, green dots illustrate the relatively few interactions for which Fisher Exact p < 0.05, but that did not pass the multiple test correction. Black and green (interchromosomal), blue (intrachromosomal).

The Fisher Exact matrix displays the expected correlation between loci linked in cis, as a diagonal line and blocks that correspond to low recombination intervals positioned around the centromeres (Fig. 3). In addition to intrachromosomal interactions, cases of strong interchromosomal ones are evident, suggesting physical linkage. For example, a region of 1Mb in chromosome 1 displays an association to the entire pericentromeric region of chromosome 8. Similar interactions are visible between chr. 4 and chr. 7 and between chr. 11 and chr. 12. These could be translocated heterochromatic blocks, which have been demonstrated in potato (Zhang *et al*. 2014; de Boer *et al*. 2015), or genomic assembly errors (see below).

An alternative approach to determining Fisher Exact probabilities, is correlation based on Pearson’s R., which while less sensitive and less interpretable could often provide equivalent information because translocation signals are strong (Methods; Suppl. Fig. 3, Suppl. Fig. 4).

**Figure 4.**
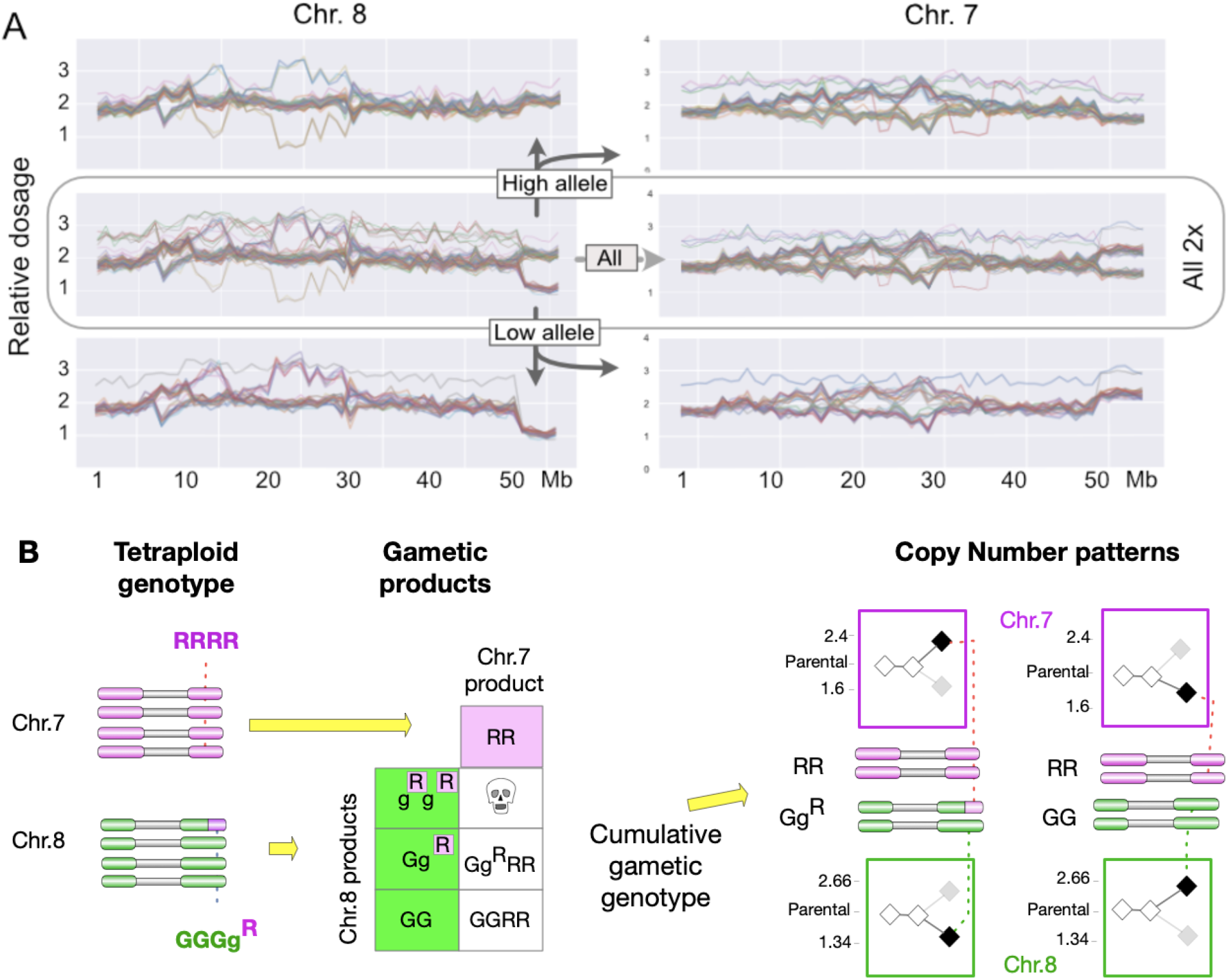
Analysis of CNV states in chromosome 7 and 8 of potato clone PI 310467. A. Complete correlation of CNV state between chr. 8 and chr. 7 in the population of dihaploids. Chr. 8 and chr. 7 relative dosages are plotted using all 2x individuals (center row) or filtered sets (top and bottom row), relative to the parental genome of PI 310467. High allele: Selection of individuals where chr. 8 distal right arm copy number > 1.5. Trisomic individuals (high dosage tracks) are removed. Low allele: Selection of individuals where chr. 8 distal right arm copy number < 1.5. The dosage of each individual was calculated by dividing standardized reads per bin by the corresponding count for PI 310467. The dosage shown is therefore not absolute, but relative to that of the maternal dosage. B. Model of Tr.8-7 in autotetraploid Alca Tarma (a.k.a. LOP868), meiotic transmission pattern of chr. 7 and chr. 8 into gametes. On the right, the genotype of dihaploids is displayed together with the resulting copy number pattern for the terminal, right arm region of chr. 7 and chr. 8. The values are relative. For example, the terminal region of 7 is present in 3 copies in the RRGg^R^ dihaploid and 5 in the tetraploid parent. The relative dosage = 3/5 * 4 = 2.4

### Translocation of a euchromatic 6Mb region between chr. 7 and 8

We noted a strong signal between the 6Mb euchromatic right arm of chr. 7 and the 5Mb right arm of chr. 8 (Fig. 3). In the dosage plots (Fig. 4), both chromosomes display distinct copy number polymorphism with absolute correlation between dosage states: higher dosage of chr. 7 was always associated with lower dosage of chr. 8. Translocation of the terminal segment of chr. 7 to chr. 8 with concurrent loss of the 5Mb terminal region of 8 (Fig. 4-B) could explain the observations, as illustrated in the inheritance and dosage model (Fig. 4-B, Suppl. Fig. 2). The dosage plots in Fig. 4 were constructed by standardizing read counts to those of parent variety PI 310467, i.e. they are relative to the control. To test the above hypothesis, we plotted raw genomic dosage of putative single copy regions corresponding to SNP (Fig. 5-C, blue track) (see Methods). We also used SNP count ratios to provide an independent measure of copy number. Fig. 5-B,C demonstrates that there are 5 copies for chr. 7 and three copies of chr. 8 in the involved regions. To provide conclusive validation for this translocation, we conducted FISH using two oligonucleotide-based chromosome painting probes specific for chr. 7 and chr. 8, respectively (Fig. 5). We observed four normal copies of chr. 7 and three normal copies of chr. 8. One additional copy of chr. 8 carried a region of chr. 7 (Fig. 5D). We concluded that the PI 310467 clone has an unbalanced translocation between chr. 7 and chr. 8. The presence of this translocation was verified by obtaining and sequencing a second, independent sample of PI 310467 from the USDA Potato Genebank (data not shown).

**Figure 5.**
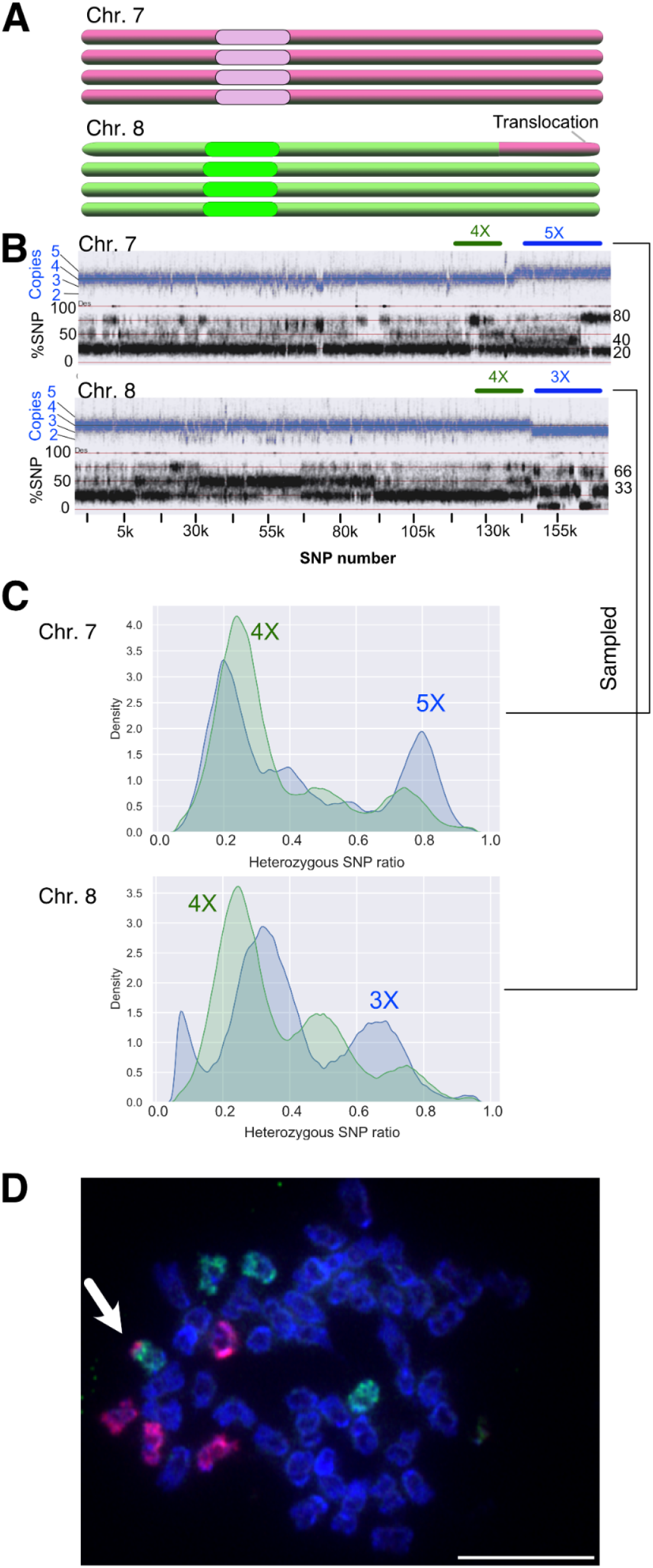
Molecular and cytological evidence for translocation 8-7. The genomic status of chr. 7 and chr. 8 is demonstrated by read coverage (copy number), SNP ratio, and oligonucleotide-based fluorescent in situ hybridization painting probes (oligo-FISH). **A**. Karyotype of chr. 7 and chr. 8 in PI 310467 according to the following evidence. **B**. The blue tracks display the DNA copies at each SNP locus along the length of the shown chromosome. The multiple black tracks illustrate the allele specific read depth ratio of SNP loci. Four DNA copies yield heterozygous SNP ratios of 25-50-75%. For example, the simplex genotype *Aaaa* corresponds to 25%. Three DNA copies yield heterozygous SNP ratios of 33-66%, five copies of 20-40-60-80%. **C**. SNP ratio analysis of chr. 7 indicating 4 and 5 copies. For chr. 8, 4 and 3 copies. **D**. Oligo-FISH painting of a mitotic metaphase cell prepared from PI 310467. The arrow points to the translocation chromosome. Red: chr. 7, Green: chr. 8.

In summary, analysis of covarying dosage states identified regions in linkage disequilibrium (i.e. genetically linked) including the unbalanced translocation between chr. 7 and chr. 8.

### Analysis of tetraploids in the BB family of potato

To probe the robustness of the method, we analyzed the tetraploid, hybrid BB progeny. These have genomes formed by the 2x egg of PI 310467 and accidental 2N (=2x) sperms from IvP48. Their dosage states derived from the additional action of alleles, two maternal and two paternal, resulting in a more complex outcome. By Fisher Exact Test we detected comparable numbers of intrachromosomal interaction (474 in the 2x group vs 525 in the 4x group), but fewer interchromosomal interactions (764 in the 2x group vs 169 in the 4x group), as indicated by the Fisher Exact matrix. For example, the matrix no longer displayed the chr. 11-12 putative translocation visible in the 2x analysis. Tr.8-7, however, was evident (Suppl. Fig. 3). We concluded that CNV resulting from duplication or deletion are more difficult to identify in a tetraploid, but events such as Tr8-7 remain distinct.

### Analysis of additional dihaploid families of potato

We asked if the method could be useful in eight comparable, but unrelated dihaploid families generated through haploid induction crosses. Seven of these dihaploid families (Amundson *et al*. 2020b) revealed the presence of translocated or misplaced heterochromatic and pericentromeric blocks, but no arm translocations (Suppl. Fig. 4, Suppl. Fig. 5). In the last dihaploid population, called LOP, an interesting arm translocation was evident that affected their tetraploid seed parent, *S. tuberosum andigena* variety Alca Tarma (Velásquez *et al*. 2007; Amundson *et al*. 2020a). Interestingly, a segment representing several Mb of euchromatin on the short arm of chr. 4 displayed a distinct dosage polymorphism that was in strong LD with the end of chr.1 (Fig. 6-A,B). Similarly to the T8-7 translocation identified in the BB population, this suggested a translocation of the euchromatic segment to the end of chr. 1, which correspondingly lost a short terminal segment. Dosage analysis in Alca Tarma demonstrated neutrality, i.e. four copies of the short arm of chr. 4. Together with the inheritance pattern observed, this suggested that the translocated chromosome T1-4 and the terminally truncated version of chr. 1 were present as a single copy in Alca Tarma (Fig. 6-C). By pooling reads from dihaploids sharing chr. 4 dosage states and rerunning the analysis with a smaller bin size, we narrowed the junction of the translocated region to a 10Kb interval between chr04.repeat.3471 and chr04.repeat.3473 (Fig. 7-A,B). This region, however, consists of N-nucleotides that could not be sequenced during the potato genome project (Fig.7-C). A model that explains the inheritance pattern and the dosage profiles of the dihaploid progeny is provided in Fig.6-C.

**Figure 6.**
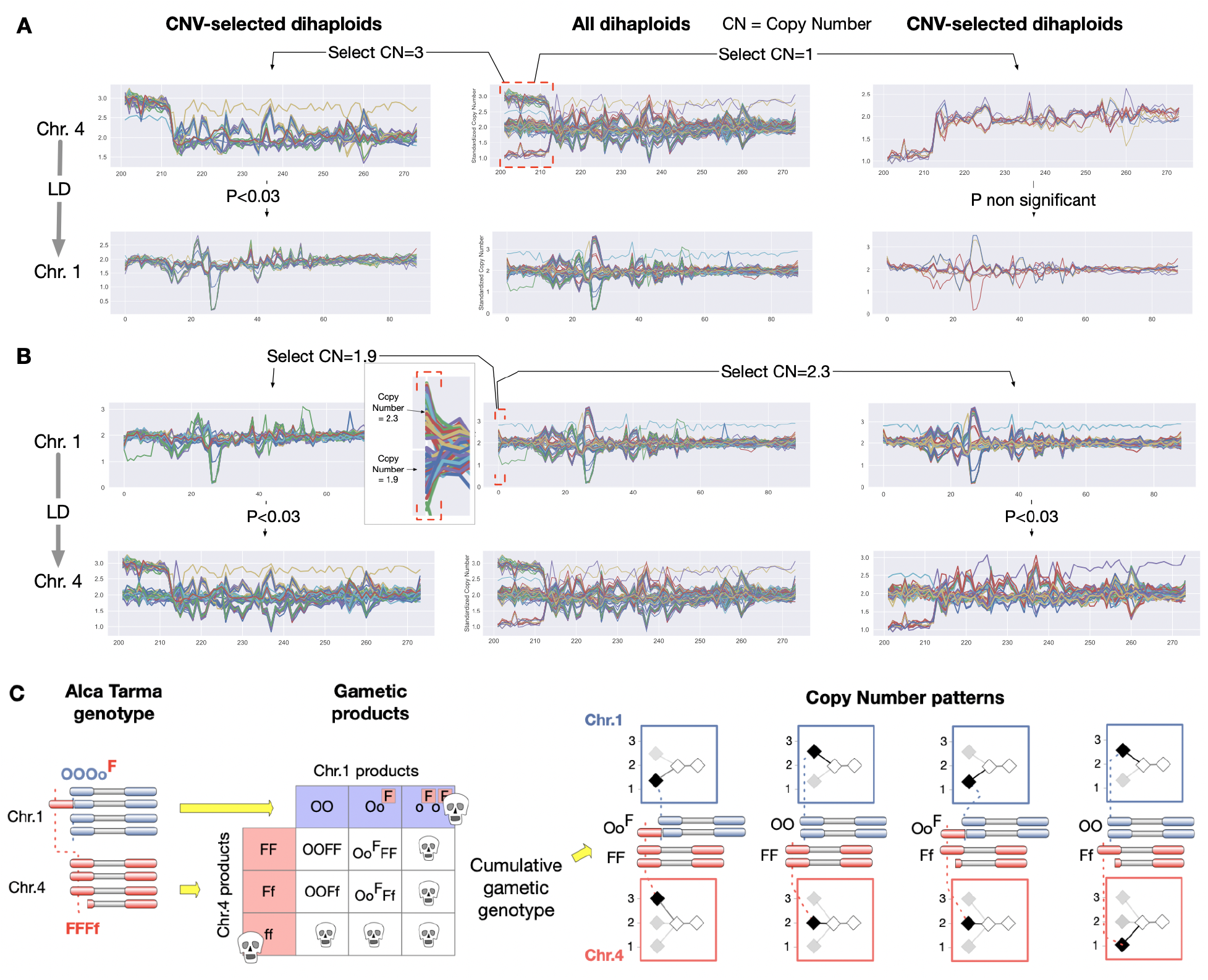
Linkage Disequilibrium (LD) consistent with translocation of a chr. 4 segment to chr. 1 in the LOP population. (A-B) Each line represents the dosage profile of an individual relative to the parental control. (A). Individuals that share either high or low cluster **Copy Number** dosage state (**CN**) on the proximal arm of chromosome 4 were selected (left, right). The chromosome 1 profiles displayed by the selected individuals confirm the LD with CN=3, the duplicated state. (B). Individuals that share either high or low CN on the proximal arm of chromosome 1 are selected. The corresponding profiles for chromosome 4 confirm LD. The left arm tip of chr. 1 is enlarged to display the two dosage states. (C). Translocation model explaining two dosage states in chr.1 and three in chr. 4. A hypothetical genotype of Alca Tarma postulates that the short arm of 4 (F) translocated to the terminal 1 region (O) producing the haplotype o^F^. Gametes carrying homozygous deletion or duplication are likely to be dead or impaired. The resulting dihaploids display CN patterns consistent with those observed. See Fig. 2 for an explanation of the different CN patterns.

**Figure 7.**
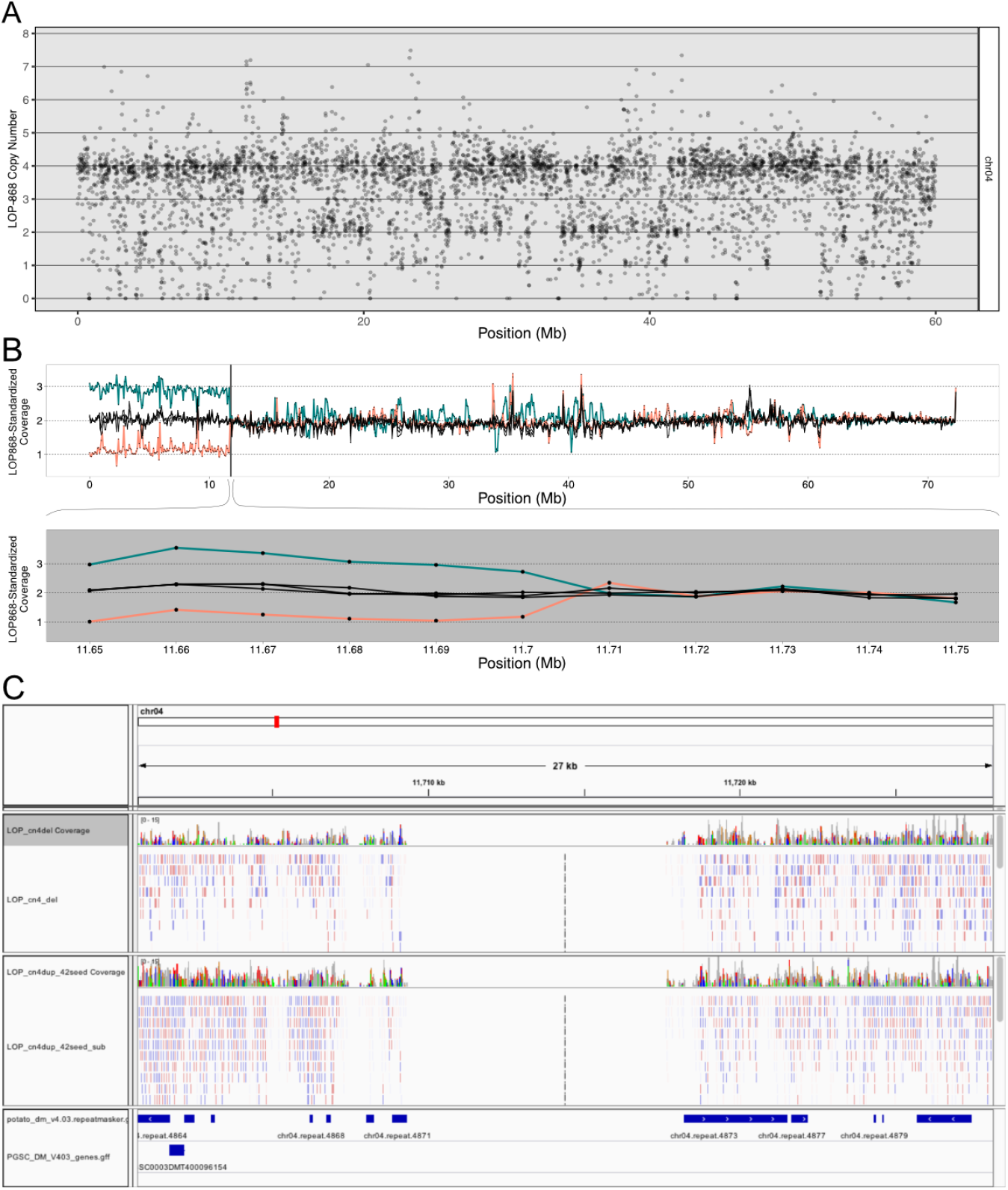
Breakpoint analysis of chromosome 4 short arm CNV. A) Relative coverage in 4x Alca Tarma over contiguous non-overlapping 1Mb bins of chromosome 4. The affected euchromatic arm is dosage neutral (4 copies). B) Top track: relative mean read coverage in different pools of dihaploids derived from Alca Tarma binned in nonoverlapping 75kb bins. The lines are colored to represent duplicated (3 copies, teal, n=28 dihaploids) or deleted (pink, n=11 dihaploids) copy number states. Shown in black are three control groups, each composed of 20 randomly selected dihaploids with neither the duplicated nor deleted state. Bottom track: the zoomed-in gray region displays higher detail (10kb bins), illustrating the change from 3 or 1 to 2 copies between 11,700,000-11,710,000 bp. C) Browser view bracketing the chromosome 4 breakpoint. The top track displays reads of pooled one-copy dihaploids, the middle track the reads of pooled 3-copy dihaploids. The breakpoint is flanked by repeats 4871 and 4873, both of which are annotated as LTR/Gypsy retrotransposons.

### Recurring translocations detected by LD-CNV result from genome misassembly

Comparison of LD in the nine dihaploid families highlighted frequent LD of certain heterochromatic regions across the populations (Suppl. Fig.3, 4, and 5). For example, a region in the left arm of chr. 1 was linked to the whole pericentromeric region of chr. 8. Similar translocations were evident for chr. 4 - 7 and chr. 11 - 12. Release of the *S. tuberosum* genome v.6.1 provided a test for the hypothesis of misassembly. A similarity dot matrix between these genome v.6.1 and genome v.404 (used here) displayed discordance for the translocated 1-8 and 4-7 regions (Suppl. Fig. 5) demonstrating that the detected misassembly was fixed independently in the update genome release.

### Detection of related CNV patterns in an A. thaliana population

We asked if CNV in LD could be identified in an unstructured natural population of *A. thaliana* (1001 Genomes Consortium 2016). We subjected 192 randomly chosen *A. thaliana* accessions that differ in geographic origin to the CNV-LD analysis (Suppl. Table 2, Suppl. Fig. 6). Overplotting CNV states displayed mostly continuous, substoichiometric variation without the frequent and distinct dosage states found in potato. We detected significant LD between chromosomes, but these affected predominantly the pericentromeric regions and were likely determined by repetitive elements because each affected loci on multiple chromosomes.

## Discussion

We developed a method that we named LD-CNV, aimed at identifying LD between copy number variable loci in the absence of genotyping information. It uses mapped reads from individuals in segregating populations. The reads are binned in genomic intervals, whose size depends on sequence coverage, but typically ranges from 0.1 to 1Mb. Read counts are normalized to the value of a reference, for example the parent, scaling the mean to overall ploidy. We identify dosage polymorphic bins by detecting variation and clustered dosage values among the tested individuals (Fig.1,2). We then test for association between dosage states of different variable bins using Fisher Exact test (Fig. 2,3). An alternative approach is to simply measure correlation using the input dosage values or the peak-centered processed values. Regardless of the statistical tool chosen, the method is simple, aiming to identify discrepancies from available genomic models. A related method, CNVmap, differs from ours by leveraging heterozygous SNP as a proxy for duplicated loci and mapping them in segregating populations (Falque *et al*. 2019). Therefore, it detects small scale duplications, but only when their sequence is divergent. The independence of LD-CNV from genotyping makes it applicable in a fully homozygous system or in any system lacking genotyping information.

We provide the software as annotated Jupyter Notebooks. These are Python-based data analysis tools that facilitate pipeline journaling, annotation, and modification for flexible analysis (Perkel 2018). In addition, plotting and display of data series from the data frames is accessible at any step of the execution (Waskom; Hunter 2007). Software dependencies can be easily installed and managed via Anaconda (“Anaconda”). Complexity and computing power of these tools are scalable, but for most analyses, such as those described here, consumer hardware is sufficient. These features should make the tool both readily usable and modifiable for ad hoc applications.

We originally developed this method to identify epistatic relationships between SV loci in potato, assuming that they would result in LD. We found, however, that the strongest signals were the result of physical LD. In our potato dihaploid progeny populations, LD-CNV analysis identified two heterochromatic blocks that were misassembled in the original reference genome and corrected in the latest one. Providing further validation, in two tetraploid seed mother accessions out of the nine tested in this analysis, we found convincing evidence of chromosomal translocations involving euchromatic arms. We detected LD between the right arms of chr. 7 and chr. 8 in the BB population (seed parent var. PI 310467), and the left arms of chr. 1 and chr. 4 in the LOP population (seed parent var. Alca Tarma). This information was combined with dosage analysis to generate translocation models. In the BB seed parent PI 310467, 5 and 3 copies, respectively, of the terminal regions of chr. 7 and chr. 8, suggested that the terminal right arm of chr. 7 translocated to chr. 8 with concurrent loss of the terminal region of chr. 8 (Fig. 6-A,B). The BB population included a tetraploid set, which enabled testing the method in a genomic scenario more challenging than diploidy. In the tetraploids, haplotypes are contributed by the tetraploid seed mother and the diploid pollen parent. Nonetheless, detection of the 8-7 translocation was still possible. The predicted rearrangement was confirmed by oligo-FISH analysis (Han *et al*. 2015; Braz *et al*. 2018) in the BB population parent. A second translocation was detected in Alca Tarma, the seed parent of LOP. In this clone, chr. 1 and chr. 4 appear copy-number neutral. This information, together with the profiles of the dihaploid progeny, supports a model in which the terminal region of chr. 4 translocated near to the end of chr. 1 (Fig. 6-C).

We wondered whether this new tool could inform us about structural variation in natural populations as well, and tested a set of 192 *A. thaliana* accessions. Comparison of the results obtained with pedigree families to this natural population shows the effect of an important feature: sibs in segregating populations form distinct dosage clusters when inheriting CNV haplotypes that are heterozygous in the parent. Large haplotypes are conserved in most individuals because recombination is rare. Large genomic bins in natural populations, however, vary continuously because of the independent behavior of multiple DNA regions and elements, hindering formation of distinct clusters. The copy number variation observed in the *A. thaliana* population corresponds most likely to repeated regions, while those observed in the potato families includes single copy sequences. The analysis should work in natural populations if the bin unit examined is very small, such as gene-size. On that scale, presence-absence of a DNA segment should cluster nicely on two dosage states. That level of granularity would require more computing power than provided by consumer hardware. If one is interested in new variation, however, LD would likely be confined to only a few bins and easily analyzed.

In conclusion, the method enables exploration of sequence datasets from segregating families in species with a reference genome to identify relationships among SV loci. Mb scale variation can be detected with as little as 0.2X coverage since one can expand bin size to increase the number of reads to a threshold of statistical confidence. Low-pass, whole-genome sequencing is a convenient approach toward genotyping that is becoming rapidly more affordable (DePristo *et al*. 2011). Once sequence reads have been mapped, the method is easily implemented without the need to identify a set of informative SNP. Knowledge of translocations, whether real or caused by genome mis-assembly, is critical to the use of a genetic population for genetic studies and for breeding because of the dramatic effect they can have on outcome and interpretation.

## Methods

### Dosage Analysis

Single end reads were aligned to the DM1-3 4.04 reference *Solanum tuberosum* genome with BWA-mem (Li 2013) and only reads with mapping quality ≥Q10 were retained. Standardized coverage values were derived by taking the fraction of mapped reads that aligned to a bin of interest for that sample, normalizing it to the corresponding fraction from the same interval in in the parent of the population, if available, or the mean of the population, and multiplying the resulting value by 2 to indicate the expected diploid state, as previously described (Henry *et al*. 2015). To mitigate mapping bias due to read type and length, paired end reads from LOP-868 were hard trimmed to 50 nt and only forward mates were used for analysis. Read depth is calculated for fixed-size non-overlapping windows, set to 250kb or 1Mb in this study. Whole chromosome aneuploids are identified as previously described (Amundson *et al*. 2020a) and withheld from analysis prior to LD matrix construction.

### Linkage Disequilibrium

The analysis and figure generating software is available at https://github.com/lcomai/cnv_mapping. Fisher’s Exact test was carried out between pairs of dosage states derived from 1Mb bins to assess linkage disequilibrium between bins. For example, assume that both Bin1 and Bin100 have three dosage states: copy number 1, 2 and 3. To test if Bin1 correlated to Bin100, the following four dihaploid sets were compared in a 2×2 contingency table: *observed in Bin1-CN1* : *observed not in Bin1-CN1, expected in Bin1-CN1* : *expected not in Bin1-CN1*, where the expectation was derived from the assumption of complete independence. Self-comparison and reciprocal comparisons were removed, and the remaining comparisons controlled at FDR = 0.05. Chromosomal bins that were engaged in at least one statistically significant correlation after these corrections were deemed in LD.

Alternative to Fisher Exact analysis of copy number dosage states, one can derive and plot a heatmap of correlation coefficients such as Pearson’s R as demonstrated in the Github notebooks (See above) and in Suppl. Fig. 3 and 4. Gating the resulting matrix to display only the strongly correlated values helps in the analysis (Suppl. Fig. 3 and 4). Measuring correlation with a utility such as *DataFrame*.*corr()* (Pandas) or *scipy*.*stats*.*pearsonr()* avoids the bin-by-bin Fisher Exact analysis, which becomes computationally intensive with increasing numbers of CNV bins. Nonetheless, we found the Fisher Exact approach manageable and clearer in its output. Categorization in peaks (dosage clusters) helped the subsequent analysis of candidate regions.

### Chromosome spread preparation

Potatoes were planted from culture tubes into a greenhouse planting mix and allowed to grow for 3-5 days. The root tips were collected and treated with iced water for 24h. Then, the root tips were fixed directly into Carnoy’s solution (3 ethanol: 1 acetic acid) and stored at −20°C until use. The root tips were squashed on the microscope slide with the same fixative solution after digestion with 2% cellulose (Sigma, USA) and 1% pectolyase (Sigma, USA) at 37°C for 2h.

### Oligo-FISH painting

We used oligonucleotide-based FISH probes (Oligo-FISH) (Han et al. 2015) to specifically paint the chromosomes 7 and 8 of potato. Probe labeling and FISH were performed following published protocols (Braz *et al*. 2018, 2020). Biotin and digoxygenin-labeled probes were detected by anti-biotin fluorescein (Vector Laboratories, Burlingame, CA, USA) and anti-digoxygenin rhodamine (Roche Diagnostics, Indianapolis, IN, USA), respectively. Chromosomes were counterstained with 4,6-diamidino-2-phenylindole in VectaShield antifade solution (Vector Laboratories). The chromosome spreads were imaged using a QImaging Retiga EXi Fast 1394 CCD camera (Teledyne Photometrics, Tucson, AZ, USA) attached to an Olympus BX51 epifluorescence microscope. Images were processed with META IMAGING SERIES 7.5 software and their final contrast was processed using Adobe PHOTOSHOP software (Adobe, San Jose, CA, USA).

## Data availability

Data availability Sequence data have been obtained from or deposited in the National Center for Biotechnology Information Sequence Read Archive with the following Bio-Project identifier: for the LOP dihaploids, PRJNA408137; BB, reads submission is pending; sequence reads data for the dihaploids of WA077, lr00014, lr00022, lr00026, 93003, C91640, C93154, are at NCBI Bioproject PRJNA699631. The raw and standardized genomic dosage values for the BB dihaploids are available at DryadData.com https://doi.org/10.25338/B88D2V. The arabidopsis sequence listed in Suppl. Table 2 was obtained from PRJNA273563 “1001 Genomes: A Catalog of Arabidopsis thaliana Genetic Variation”.

## Acknowledgments

This work was supported by: the National Science Foundation Plant Genome Integrative Organismal Systems (IOS) Grants 1444612 (Rapid and Targeted Introgression of Traits via Genome Elimination) and 1956429 (RESEARCH-PGR: Variants and Recombinants without Meiosis) to L.C. and I.M.H; a grant from the Innovative Genome Institute at UC Berkeley to LC, and the United States - Israel Binational Agricultural Research and Development Funds IS-5038-17C and IS-5317-20C to J.J.

## Author contributions

L.C. designed experiments with input from K.R.A. and I.M.H.. L.C., K.R.A., and B.O performed genomic experiments. XZ, GTB, and JJ performed cytological experiments. L.C. analyzed data and wrote the manuscript with input from all authors.

## Supplemental Figures and Tables

**Supplemental Fig. 1.**
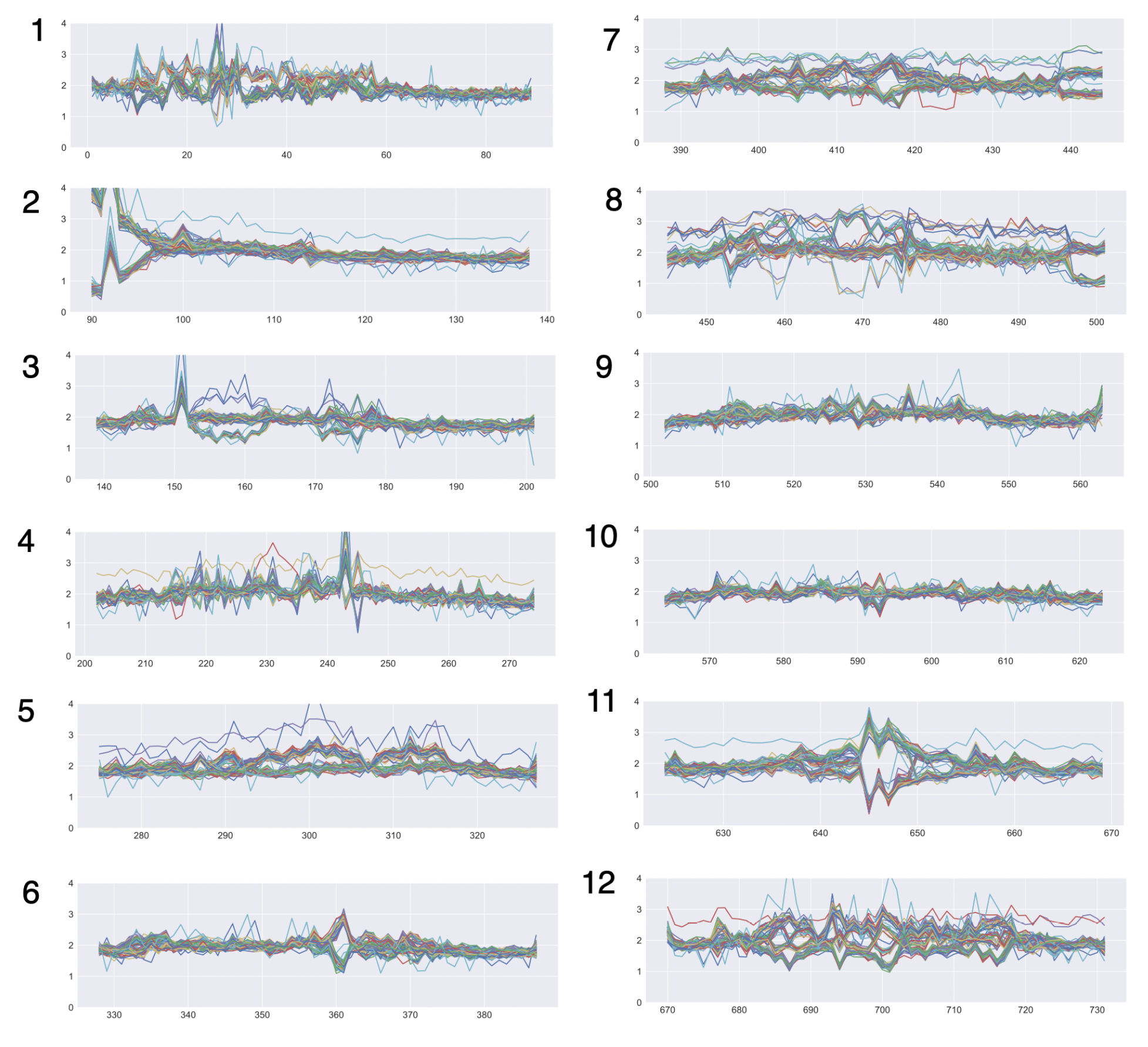
CNV patterns in BB, the dihaploid population of potato cv. PI 310467. The dosage tracks of 84 2x individuals produced by crossing cv. PI 310467 to haploid inducer IvP48.

**Supplemental Fig. 2.**
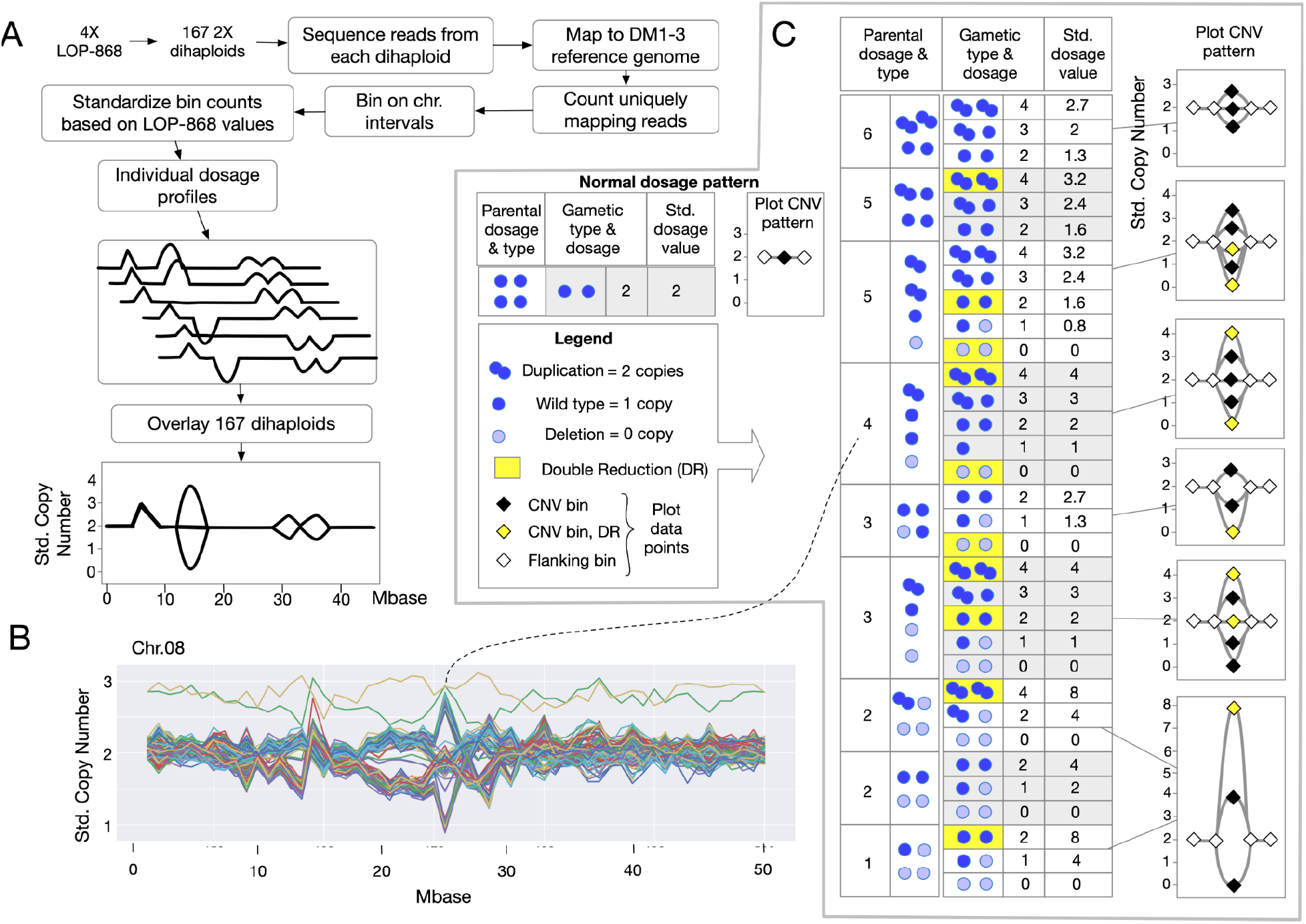
Analysis of CNV. A. Overview of method to produce the dosage plot using a dihaploid family produced from an autotetraploid parent (LOP868 is the CIP accession name for Alca Tarma, the parent of the LOP population). B. Example of dosage plot resulting from the plotting of dihaploid profiles. Each color represents an individual. The two outliers are trisomics for chr. 8. C. Dosage variation in the autotetraploid parent and resulting segregation patterns in the dihaploids, which represent the maternal gametes. The relative dosage is calculated from the formula: (CN^2x^/CN^4x^) * 4, where *CN* is the copy number and *nx* the ploidy of the individual. Dark blue: present allele. Light blue: absent allele (deletion). Overlapping circles represent a duplicated allele inherited as a unit. Yellow cells represent gametes formed by double reduction. Double reduction occurs at low frequency and is dependent on the recombination frequency between the locus of interest and the centromere.

**Supplemental Fig. 3.**
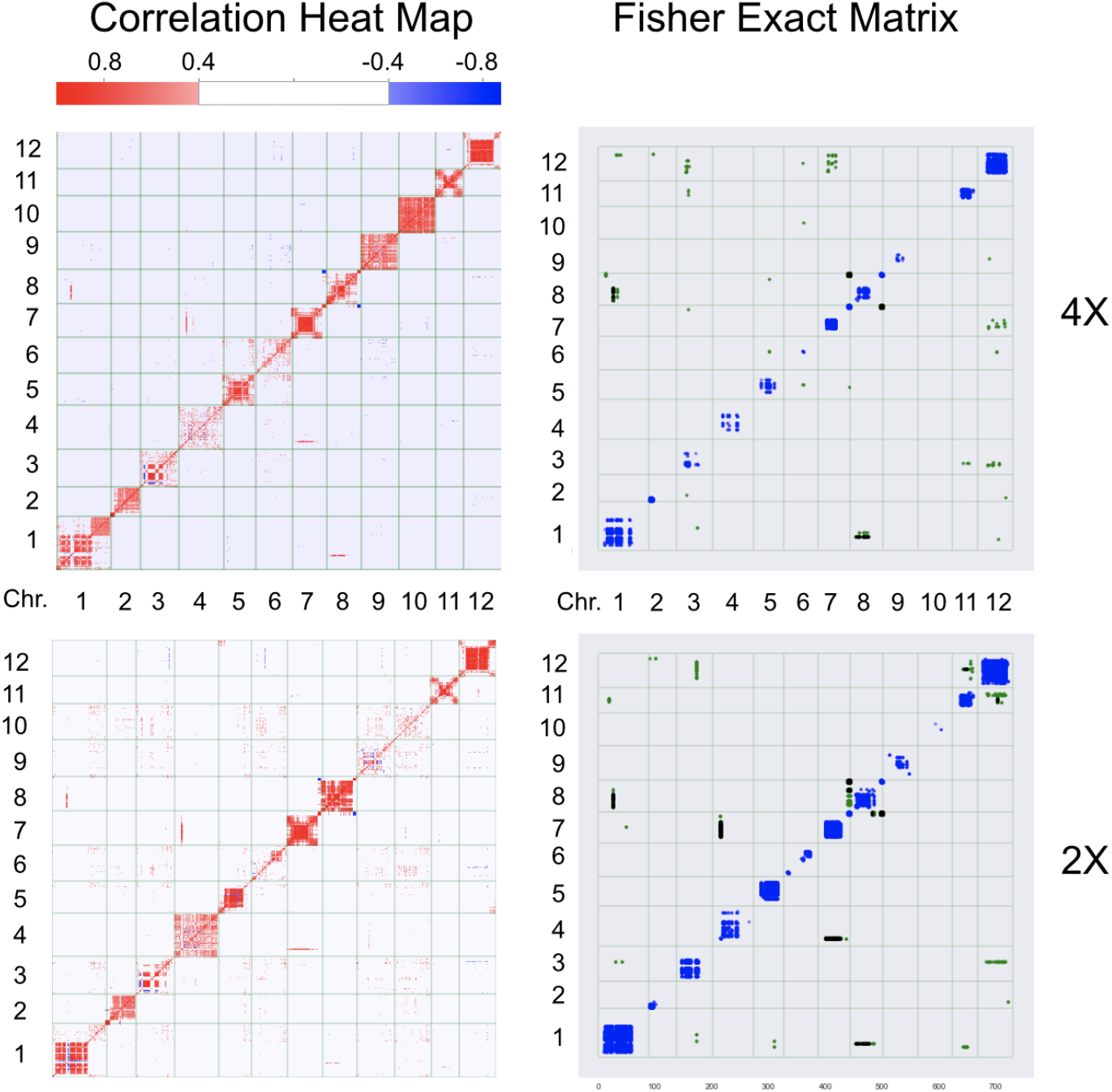
Effect of ploidy and statistical method on LD analysis. 4X analyses used the tetraploids of the BB population. 2X analyses used the diploids. The use of the Pearson’s Correlation matrix and the implementation of the Fisher Exact matrix are presented in Methods and are available at https://github.com/lcomai/cnv_mapping.

**Supplemental Fig. 4.**
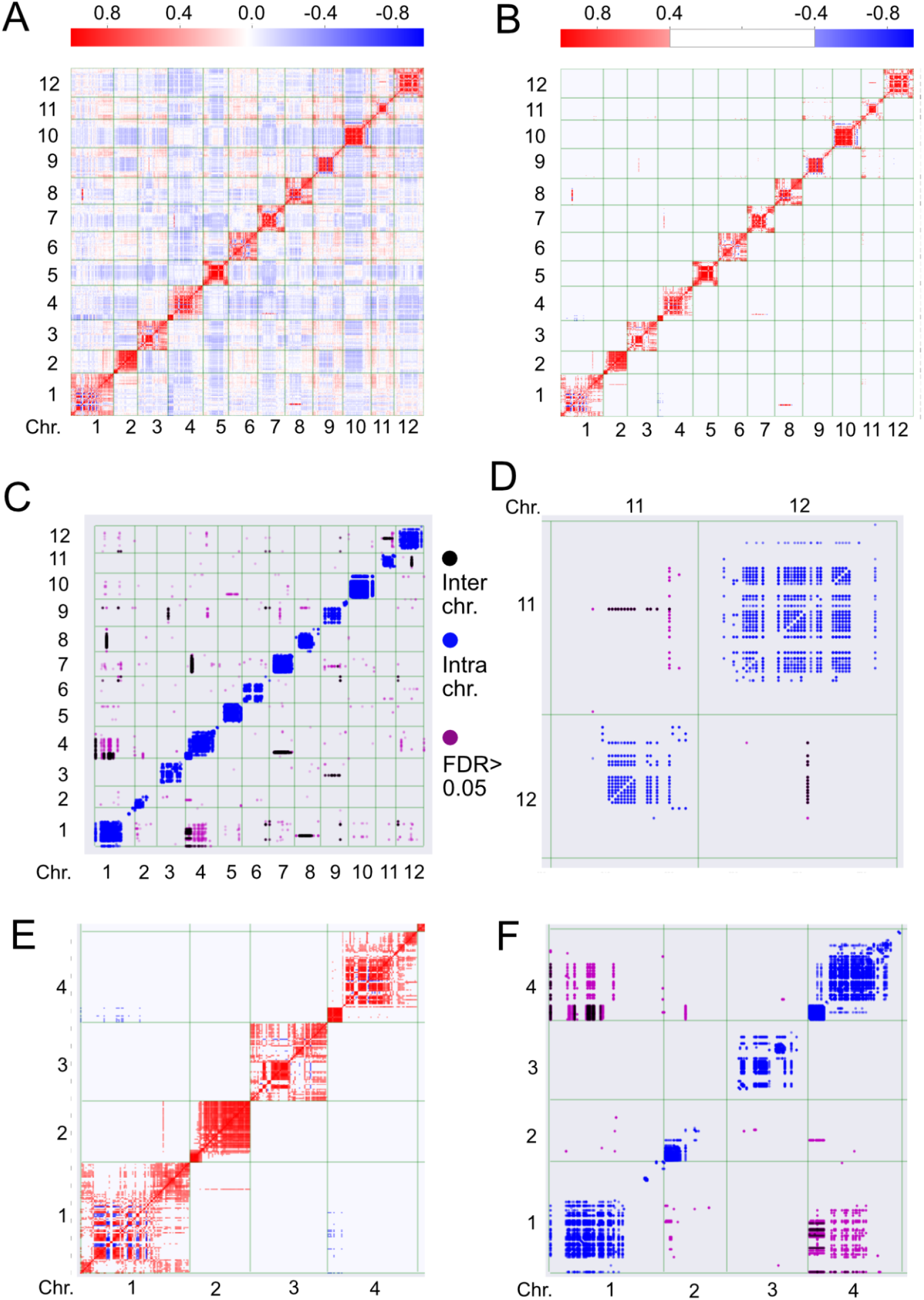
Analysis of the LOP dihaploid family and comparison of Pearson’s Correlation and Fisher Exact test for detection of correlated CNV sites. A. Raw correlation heatmap is less specific and more difficult to interpret than the Fisher Exact matrix. B. Improved heatmap display by filtering low correlation signals. **C**. Clear demarcation of candidate correlated regions by cluster analysis and Fisher Exact test. A major signal is visible for chromosome 1 and 4. D. Fisher Exact identifies candidate regions in chr. 11 and 12. This appears to be the same signal displayed by the BB population, suggesting that two different cultivated potatoes share polymorphism. E, F. Side by side comparison of translocation signals in filtered correlation and Fishes Exact matrices.

**Supplemental Fig. 5.**
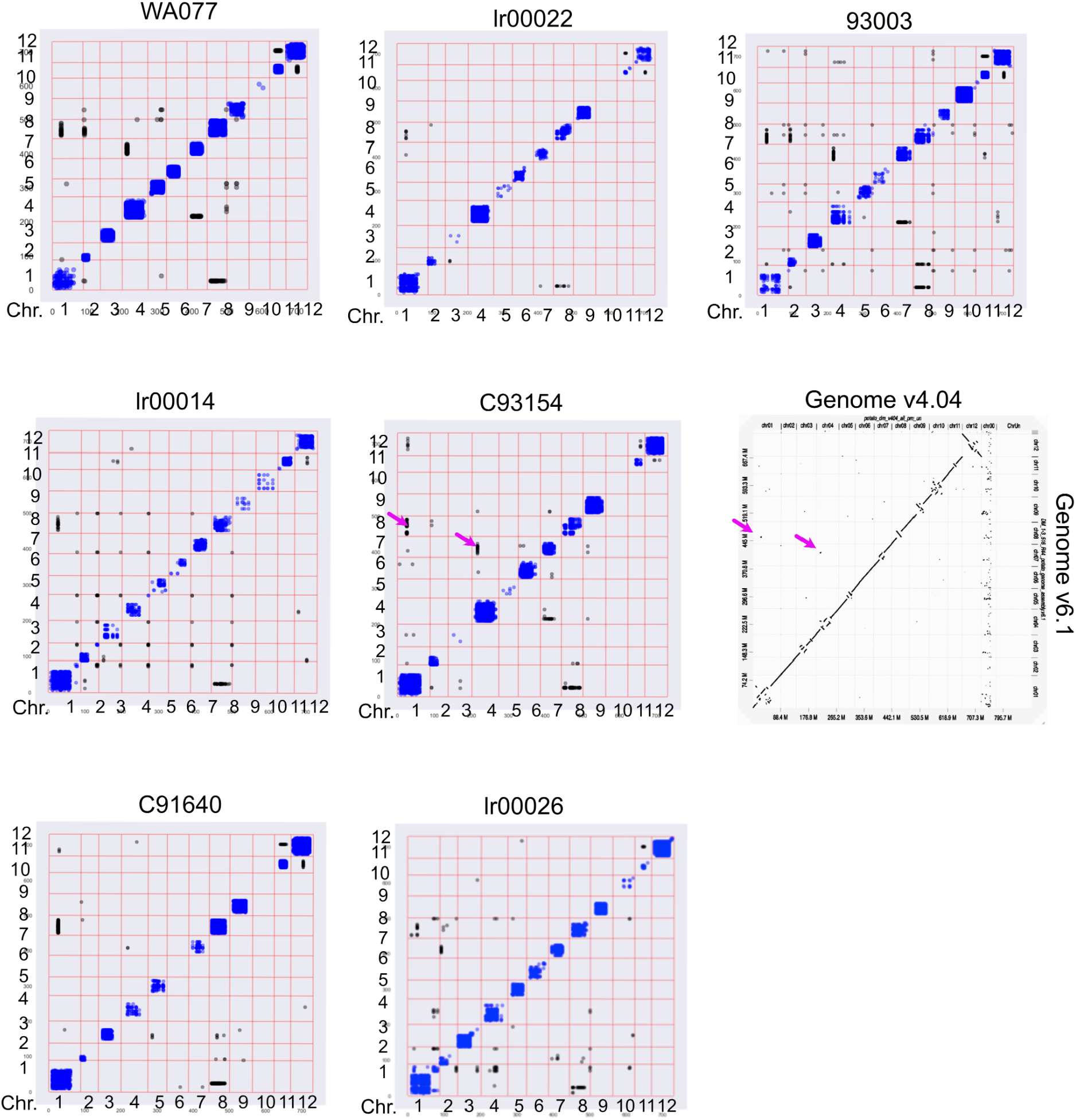
Fisher Exact LD matrices from seven potato populations. The name of each dihaploid family seed parent is shown on top of each matrix. Blue: intrachromosomal LD. Black: interchromosomal LD. In all cases, the interchromosomal signals are consistent with pericentromeric translocation or misassembly during construction of the reference genome. The 8th matrix is a dot matrix comparison (Cabanettes and Klopp 2018) of the genome assembly version used in this work (4.04) to a more recent one (6.1). The magenta arrows point to regions in chromosome 8 vs 1 and 7 vs 4 that were misassembled in 4.04 and appear “translocated” in multiple population analyses. The absence of the signal in some populations could be due to the lack of segregating CNV in those regions.

**Supplemental Fig. 6.**
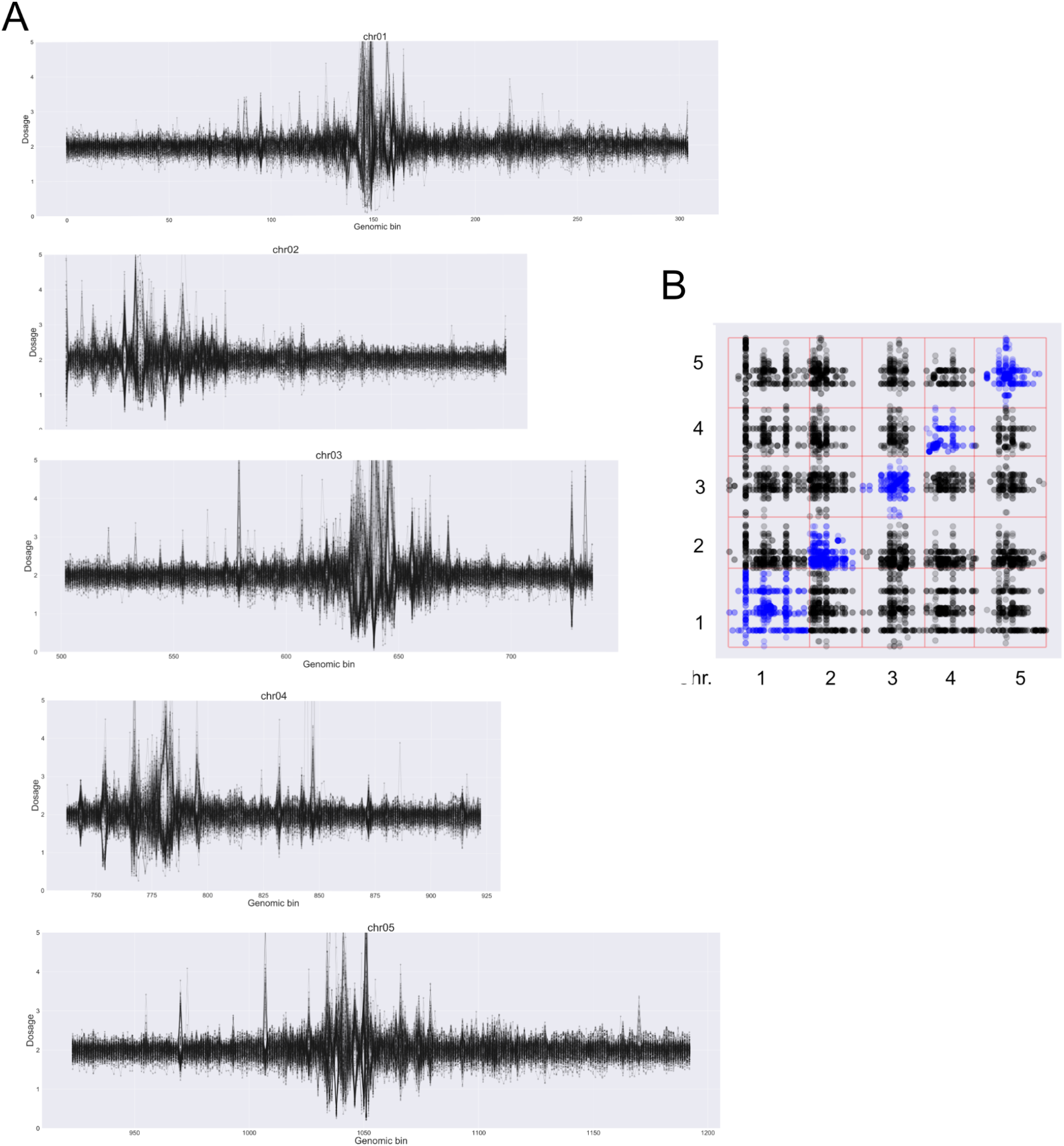
Analysis of CNV in Arabidopsis natural accessions. A. Chromosomal dosage profiles (chr. 1 to chr. 5, top to bottom) for the 192 *A. thaliana* accessions in Supplemental Table 1. B. Linkage Disequilibrium Matrix of Fisher Exact dosage correlation analysis for bins with FDR = 0.05 and for which the dosage difference between peaks is at least 1. Black: interchromosomal, blue: intrachromosomal. The interchromosomal signals demonstrate LD common to most pericentromeric regions and therefore likely caused by repeats.

**Supplemental Table 1.** List of potato accessions and sequencing library deposit

| Population Name | Tetraploid parent | Haploid inducer | Project number in NCBI read sequence database |
| --- | --- | --- | --- |
| LOP | Alca Tarma, a.k.a LOP 868 | PL-4, IvP-101 | PRJNA408137 |
| BB | PI 310467, cv. Desiree in germplasm collection | IvP-48 | Raw sequence in progress. Binned dosage values are at https://doi.org/10.25338/B88D2V |
| WA077 | WA077 (CIP397077.16 or cv. Alliance/Sarnav) | PL-4, IvP-101 | PRJNA699631 |
| lr00014 | lr00014 (CIP300056.33) |  |  |
| lr00022 | lr00022 (CIP300072.1) |  |  |
| lr00026 | lr00026 (CIP300093.14) |  |  |
| 93003 | 93003 (CIP390637.1) |  |  |
| C91640 | C91640 (CIP388615.22) |  |  |
| C93154 | C93154 (CIP392820.1 or cv. BARI Alu-73) |  |  |

**Supplemental Table 2.** List of Arabidopsis accessions, SRA number and sequencing library characteristics

| **Accession + SRR (run name in SRA db at NCBI PRJNA273563)** | **Reads** | **Reads/MB** |
| --- | --- | --- |
| admixed-GMI-10_SRR1946059 | 59916068 | 500686.84 |
| admixed-GMI-11_SRR1945728 | 24022752 | 200745.41 |
| admixed-GMI-12_SRR1945997 | 29659042 | 247844.9 |
| admixed-GMI-13_SRR1946061 | 33793413 | 282393.65 |
| admixed-GMI-14_SRR1946077 | 34644790 | 289508.16 |
| admixed-GMI-15_SRR1945983 | 19010462 | 158860.36 |
| admixed-GMI-1_SRR1945626 | 19800036 | 165458.41 |
| admixed-GMI-2_SRR1946071 | 28704624 | 239869.34 |
| admixed-GMI-3_SRR1945491 | 40568849 | 339012.38 |
| admixed-GMI-4_SRR1946022 | 36515150 | 305137.77 |
| admixed-GMI-5_SRR1945490 | 16039451 | 134033.2 |
| admixed-GMI-6_SRR1945623 | 19421444 | 162294.72 |
| admixed-GMI-7_SRR1946069 | 21392755 | 178767.92 |
| admixed-GMI-8_SRR1946070 | 30120943 | 251704.77 |
| admixed-GMI-9_SRR1945638 | 32841499 | 274439.01 |
| admixed-Monsanto-10_SRR1945537 | 9055319 | 75670.5 |
| admixed-Monsanto-11_SRR1946183 | 16810601 | 140477.29 |
| admixed-Monsanto-12_SRR1946126 | 17673420 | 147687.41 |
| admixed-Monsanto-13_SRR1946184 | 21228594 | 177396.12 |
| admixed-Monsanto-14_SRR1946471 | 13767340 | 115046.37 |
| admixed-Monsanto-15_SRR1946400 | 12540265 | 104792.35 |
| admixed-Monsanto-1_SRR1945940 | 10699758 | 89412.21 |
| admixed-Monsanto-2_SRR1945519 | 15019671 | 125511.43 |
| admixed-Monsanto-3_SRR1946108 | 17166146 | 143448.39 |
| admixed-Monsanto-4_SRR1946149 | 15577834 | 130175.71 |
| admixed-Monsanto-5_SRR1946418 | 5990357 | 50058.24 |
| admixed-Monsanto-6_SRR1946135 | 17295878 | 144532.49 |
| admixed-Monsanto-7_SRR1945556 | 9700463 | 81061.63 |
| admixed-Monsanto-8_SRR1946479 | 17924451 | 149785.14 |
| admixed-Monsanto-9_SRR1946438 | 13674684 | 114272.09 |
| asia-GMI-1_SRR1946559 | 28507994 | 238226.21 |
| asia-GMI-2_SRR1946560 | 55378823 | 462771.49 |
| asia-GMI-3_SRR1946561 | 41916903 | 350277.36 |
| asia-GMI-4_SRR1946558 | 25864327 | 216134.48 |
| asia-GMI-5_SRR1946555 | 27246541 | 227684.91 |
| asia-GMI-6_SRR1946566 | 35257170 | 294625.49 |
| asia-GMI-7_SRR1946557 | 24278378 | 202881.54 |
| asia-GMI-8_SRR1946565 | 41394961 | 345915.76 |
| asia-GMI-9_SRR1946556 | 27748807 | 231882.08 |
| asia-Monsanto-10_SRR1946049 | 30564177 | 255408.64 |
| asia-Monsanto-11_SRR1946212 | 12655516 | 105755.44 |
| asia-Monsanto-12_SRR1946225 | 12970324 | 108386.13 |
| asia-Monsanto-13_SRR1946218 | 9342590 | 78071.08 |
| asia-Monsanto-14_SRR1946312 | 9520464 | 79557.47 |
| asia-Monsanto-15_SRR1946203 | 13293030 | 111082.81 |
| asia-Monsanto-1_SRR1946229 | 11167396 | 93320.01 |
| asia-Monsanto-2_SRR1946214 | 9013359 | 75319.87 |
| asia-Monsanto-3_SRR1946219 | 8855134 | 73997.66 |
| asia-Monsanto-4_SRR1946224 | 9790332 | 81812.62 |
| asia-Monsanto-5_SRR1946199 | 12179493 | 101777.57 |
| asia-Monsanto-6_SRR1946201 | 11459167 | 95758.19 |
| asia-Monsanto-7_SRR1946221 | 12692203 | 106062.02 |
| asia-Monsanto-8_SRR1946050 | 23325425 | 194918.22 |
| asia-Monsanto-9_SRR1946216 | 11455734 | 95729.5 |
| central_europe-GMI-1_SRR1946008 | 19335113 | 161573.3 |
| central_europe-GMI-2_SRR1946569 | 40264985 | 336473.15 |
| central_europe-GMI-3_SRR1946563 | 69280709 | 578942.19 |
| central_europe-GMI-4_SRR1946562 | 63443436 | 530163.19 |
| central_europe-GMI-5_SRR1945724 | 20054005 | 167580.7 |
| central_europe-GMI-6_SRR1946564 | 68211731 | 570009.3 |
| central_europe-GMI-7_SRR1946568 | 36714388 | 306802.69 |
| central_europe-GMI-8_SRR1946567 | 44576218 | 372499.84 |
| central_europe-Monsanto-10_SRR1946230 | 12765831 | 106677.29 |
| central_europe-Monsanto-11_SRR1946268 | 19014619 | 158895.1 |
| central_europe-Monsanto-12_SRR1945887 | 11104130 | 92791.33 |
| central_europe-Monsanto-13_SRR1946343 | 20579298 | 171970.29 |
| central_europe-Monsanto-14_SRR1946465 | 11770417 | 98359.14 |
| central_europe-Monsanto-15_SRR1946247 | 10763715 | 89946.66 |
| central_europe-Monsanto-1_SRR1946249 | 9821512 | 82073.17 |
| central_europe-Monsanto-2_SRR1946252 | 13265534 | 110853.04 |
| central_europe-Monsanto-3_SRR1945443 | 16016473 | 133841.18 |
| central_europe-Monsanto-4_SRR1946263 | 15117094 | 126325.55 |
| central_europe-Monsanto-5_SRR1946254 | 14385342 | 120210.68 |
| central_europe-Monsanto-6_SRR1946272 | 11963830 | 99975.39 |
| central_europe-Monsanto-7_SRR1946266 | 20927166 | 174877.24 |
| central_europe-Monsanto-8_SRR1946363 | 22051503 | 184272.73 |
| central_europe-Monsanto-9_SRR1946369 | 21010693 | 175575.23 |
| germany-GMI-10_SRR1946094 | 64112581 | 535754.88 |
| germany-GMI-11_SRR1946103 | 20588686 | 172048.74 |
| germany-GMI-12_SRR1945757 | 14245174 | 119039.37 |
| germany-GMI-1_SRR1946009 | 17541288 | 146583.25 |
| germany-GMI-2_SRR1945729 | 79383009 | 663361.76 |
| germany-GMI-3_SRR1945696 | 25408557 | 212325.85 |
| germany-GMI-4_SRR1945793 | 28859058 | 241159.86 |
| germany-GMI-5_SRR1945727 | 10380604 | 86745.21 |
| germany-GMI-6_SRR1945726 | 23114360 | 193154.46 |
| germany-GMI-7_SRR1945943 | 17913268 | 149691.69 |
| germany-GMI-8_SRR1945944 | 10227695 | 85467.43 |
| germany-GMI-9_SRR1945730 | 57664135 | 481868.63 |
| germany-Monsanto-10_SRR1945873 | 9570405 | 79974.81 |
| germany-Monsanto-11_SRR1945468 | 15082766 | 126038.69 |
| germany-Monsanto-12_SRR1945575 | 17094637 | 142850.83 |
| germany-Monsanto-13_SRR1945960 | 8887338 | 74266.78 |
| germany-Monsanto-14_SRR1945507 | 23900073 | 199720.25 |
| germany-Monsanto-15_SRR1945513 | 15828270 | 132268.47 |
| germany-Monsanto-1_SRR1945515 | 14431836 | 120599.21 |
| germany-Monsanto-2_SRR1945474 | 10955767 | 91551.54 |
| germany-Monsanto-3_SRR1945473 | 22297592 | 186329.17 |
| germany-Monsanto-4_SRR1945961 | 9722146 | 81242.82 |
| germany-Monsanto-5_SRR1945538 | 14253405 | 119108.16 |
| germany-Monsanto-6_SRR1946469 | 15668800 | 130935.86 |
| germany-Monsanto-7_SRR1945583 | 22458306 | 187672.17 |
| germany-Monsanto-8_SRR1945531 | 19783897 | 165323.55 |
| germany-Monsanto-9_SRR1945554 | 9266572 | 77435.83 |
| italy_balkan_caucasus-Monsanto-10_SRR1946282 | 7402735 | 61860.74 |
| italy_balkan_caucasus-Monsanto-11_SRR1946297 | 19763662 | 165154.45 |
| italy_balkan_caucasus-Monsanto-12_SRR1946240 | 9828609 | 82132.48 |
| italy_balkan_caucasus-Monsanto-13_SRR1946032 | 18279195 | 152749.55 |
| italy_balkan_caucasus-Monsanto-14_SRR1946321 | 10435351 | 87202.7 |
| italy_balkan_caucasus-Monsanto-15_SRR1946323 | 21265077 | 177700.98 |
| italy_balkan_caucasus-Monsanto-1_SRR1946288 | 11969392 | 100021.87 |
| italy_balkan_caucasus-Monsanto-2_SRR1946046 | 16701439 | 139565.08 |
| italy_balkan_caucasus-Monsanto-3_SRR1946301 | 9280705 | 77553.94 |
| italy_balkan_caucasus-Monsanto-4_SRR1946023 | 13498714 | 112801.6 |
| italy_balkan_caucasus-Monsanto-5_SRR1946291 | 11484184 | 95967.24 |
| italy_balkan_caucasus-Monsanto-6_SRR1946036 | 31954866 | 267029.89 |
| italy_balkan_caucasus-Monsanto-7_SRR1946292 | 14538360 | 121489.37 |
| italy_balkan_caucasus-Monsanto-8_SRR1946025 | 19477965 | 162767.04 |
| italy_balkan_caucasus-Monsanto-9_SRR1946295 | 11013689 | 92035.57 |
| north_sweden-GMI-10_SRR1945690 | 35646246 | 297876.8 |
| north_sweden-GMI-11_SRR1945716 | 13063824 | 109167.46 |
| north_sweden-GMI-12_SRR1946090 | 37244466 | 311232.27 |
| north_sweden-GMI-13_SRR1946007 | 18173335 | 151864.93 |
| north_sweden-GMI-14_SRR1945632 | 21309495 | 178072.16 |
| north_sweden-GMI-15_SRR1945630 | 26677852 | 222932.68 |
| north_sweden-GMI-1_SRR1945604 | 16929373 | 141469.8 |
| north_sweden-GMI-2_SRR1945715 | 16638157 | 139036.26 |
| north_sweden-GMI-3_SRR1945697 | 34935477 | 291937.28 |
| north_sweden-GMI-4_SRR1945694 | 26013634 | 217382.16 |
| north_sweden-GMI-5_SRR1945607 | 38118179 | 318533.43 |
| north_sweden-GMI-6_SRR1945605 | 58226536 | 486568.32 |
| north_sweden-GMI-7_SRR1945711 | 36444439 | 304546.87 |
| north_sweden-GMI-8_SRR1945691 | 24890971 | 208000.66 |
| north_sweden-GMI-9_SRR1946064 | 17125390 | 143107.81 |
| relict-Monsanto-10_SRR1946427 | 11670262 | 97522.2 |
| relict-Monsanto-11_SRR1946195 | 11254133 | 94044.83 |
| relict-Monsanto-12_SRR1946393 | 13088955 | 109377.46 |
| relict-Monsanto-13_SRR1946152 | 18967479 | 158501.18 |
| relict-Monsanto-14_SRR1946443 | 16060723 | 134210.95 |
| relict-Monsanto-15_SRR1946398 | 17903397 | 149609.21 |
| relict-Monsanto-1_SRR1946147 | 20211669 | 168898.21 |
| relict-Monsanto-2_SRR1946143 | 14356021 | 119965.66 |
| relict-Monsanto-3_SRR1946168 | 16425998 | 137263.36 |
| relict-Monsanto-4_SRR1946140 | 17347996 | 144968.01 |
| relict-Monsanto-5_SRR1946192 | 20661996 | 172661.36 |
| relict-Monsanto-6_SRR1946190 | 13805415 | 115364.54 |
| relict-Monsanto-7_SRR1946436 | 18121950 | 151435.54 |
| relict-Monsanto-8_SRR1946132 | 16429270 | 137290.71 |
| relict-Monsanto-9_SRR1946458 | 9246205 | 77265.64 |
| south_sweden-GMI-10_SRR1946101 | 40376693 | 337406.64 |
| south_sweden-GMI-11_SRR1945636 | 15148525 | 126588.2 |
| south_sweden-GMI-12_SRR1945648 | 39477578 | 329893.21 |
| south_sweden-GMI-13_SRR1945698 | 19662788 | 164311.5 |
| south_sweden-GMI-14_SRR1946100 | 38782621 | 324085.82 |
| south_sweden-GMI-15_SRR1945660 | 27492214 | 229737.87 |
| south_sweden-GMI-1_SRR1945633 | 35549994 | 297072.47 |
| south_sweden-GMI-2_SRR1945637 | 38539759 | 322056.35 |
| south_sweden-GMI-3_SRR1945683 | 35359557 | 295481.09 |
| south_sweden-GMI-4_SRR1945996 | 28208636 | 235724.63 |
| south_sweden-GMI-5_SRR1945964 | 15576292 | 130162.82 |
| south_sweden-GMI-6_SRR1945477 | 19110039 | 159692.47 |
| south_sweden-GMI-7_SRR1945707 | 30146186 | 251915.71 |
| south_sweden-GMI-8_SRR1946087 | 39000378 | 325905.5 |
| south_sweden-GMI-9_SRR1946092 | 23810485 | 198971.61 |
| south_sweden-Monsanto-1_SRR1945906 | 11470851 | 95855.83 |
| spain-Monsanto-10_SRR1946155 | 15160643 | 126689.46 |
| spain-Monsanto-11_SRR1946151 | 13526890 | 113037.05 |
| spain-Monsanto-12_SRR1946402 | 14183058 | 118520.3 |
| spain-Monsanto-13_SRR1946435 | 16866069 | 140940.8 |
| spain-Monsanto-14_SRR1946412 | 12964849 | 108340.38 |
| spain-Monsanto-15_SRR1946451 | 16753696 | 140001.76 |
| spain-Monsanto-1_SRR1946437 | 13150132 | 109888.69 |
| spain-Monsanto-2_SRR1946381 | 12589966 | 105207.68 |
| spain-Monsanto-3_SRR1946404 | 8464114 | 70730.12 |
| spain-Monsanto-4_SRR1946453 | 15113990 | 126299.61 |
| spain-Monsanto-5_SRR1946426 | 14461779 | 120849.43 |
| spain-Monsanto-6_SRR1946154 | 17281639 | 144413.5 |
| spain-Monsanto-7_SRR1946133 | 20868196 | 174384.46 |
| spain-Monsanto-8_SRR1946167 | 18623040 | 155622.88 |
| spain-Monsanto-9_SRR1946454 | 18006844 | 150473.66 |
| western_europe-GMI-1_SRR1945657 | 26646916 | 222674.16 |
| western_europe-Monsanto-10_SRR1946389 | 13761980 | 115001.58 |
| western_europe-Monsanto-11_SRR1945553 | 7896785 | 65989.25 |
| western_europe-Monsanto-12_SRR1946480 | 11433818 | 95546.36 |
| western_europe-Monsanto-13_SRR1945456 | 21387945 | 178727.73 |
| western_europe-Monsanto-14_SRR1946407 | 19483841 | 162816.14 |
| western_europe-Monsanto-15_SRR1945552 | 16259172 | 135869.29 |
| western_europe-Monsanto-1_SRR1946464 | 7161758 | 59847.02 |
| western_europe-Monsanto-2_SRR1946372 | 15399476 | 128685.26 |
| western_europe-Monsanto-3_SRR1945568 | 21045538 | 175866.41 |
| western_europe-Monsanto-4_SRR1945891 | 19333214 | 161557.43 |
| western_europe-Monsanto-5_SRR1946055 | 12240368 | 102286.27 |
| western_europe-Monsanto-6_SRR1946116 | 11464228 | 95800.48 |
| western_europe-Monsanto-7_SRR1946460 | 10741251 | 89758.95 |
| western_europe-Monsanto-8_SRR1945557 | 14674892 | 122630.3 |
| western_europe-Monsanto-9_SRR1945438 | 18291432 | 152851.81 |

## Notes

https://github.com/lcomai/cnv_mapping

### Competing Interest Statement

The authors have declared no competing interest.

### Summary of Updates

minor fixes

